# Mechanical communication-induced cell directional migration and branching connections mediated by calcium channels, integrin β1 and N-cadherin

**DOI:** 10.1101/2022.05.02.490256

**Authors:** Mingxing Ouyang, Yiming Zhu, Jiajia Wang, Qingyu Zhang, Bing Bu, Jia Guo, Linhong Deng

## Abstract

Cell-cell mechanical communications at large spatial scale (above hundreds of micrometers) have been increasingly recognized in recent decade, which shows importance in tissue-level assembly and morphodynamics. The involved mechanosensing mechanism and resulted physiological functions are still to be fully understood. Recent work showed that traction force sensation in the matrix induces cell communications for self-assembly. Here, based on the experimental model of cell directional migration on Matrigel hydrogel containing 0.5 mg/ml type I collagen, we studied the mechano-responsive pathways for cell distant communications. Airway smooth muscle (ASM) cells assembled network structure on the hydrogel, whereas stayed isolated individually when cultured on glass without force transmission. Cell directional migration, or network assembly was significantly attenuated by inhibited actomyosin activity, or inhibition of inositol 1,4,5-trisphosphate receptor (IP_3_R) calcium channel or SERCA pump on endoplasmic reticulum (ER) membrane, or L-type calcium channel on the plasma membrane. Inhibition of integrin β1 with siRNA knockdown reduced cell directional migration and branching assembly, whereas inhibition of cell junctional N-cadherin with siRNA had little effect on distant attractions but blocked branching assembly. Our work demonstrated that the ER calcium channels and integrin are mechanosensing signals for cell mechanical communications regulated by actomyosin activity, while N-cadherin is responsible for traction force-induced cell stable connections in the assembly.

## Introduction

Cell-cell mechanical communications in distance have been much documented in recent decade, which has shown remote mechanosensitive responses among cells[1; 2]. In tracking the development of this discovery back, biologists had observed the outgrowth of separated neuronal tissues towards each other in early last century [3; 4]. In 1981, Harris et al. reported that traction force from fibroblast tissue explants induced fibrillary morphogenesis of type I collagen (COL), which separation reached a distance of 1.5 cm[5]. Over ten years ago, single-cell studies reported mechanical interactions within neighboring cells[6; 7]. Later, studies established long-range force-induced communications among mammary cell clusters, which causes COL fiber modeling and helps assembly of tissue patterns[8; 9; 10]. Therefore, the time line of early developments crosses near one century, although some these studies might have been done without knowledge of each other.

Within recent decade, more works have demonstrated cell long-range mechanical communications under different scenarios, such as synchronized beating of cardiomyocytes[11; 12], collective migration of epithelial sheet[13], long-range aligned pattern of fibroblasts[14], myofibroblast-fibroblast cross-talk in fibrosis[15], and remote attraction of macrophages by fibroblasts[16]. At the same time, the relevant mechanism of long-range force transmission and COL fibrillary modeling has attracted research enthusiasms based on experimental data and computational simulations[17; 18; 19]. The property of nonlinear elastic extracellular matrix (ECM) from tissue can facilitate distant force transmission[20], and cell-generated stresses are capable of buckling COL filaments in ECM at large spatial extent[21; 22]. Our recent data has shown that motile cells can actively recruit environmental COL to assemble filaments for tissue-level assembly[23].

These observations also lead to an interesting topic how cells mechanosense each other in distance or at large-spatial scales. Some progresses have emerged in recent years, for example, mechanosensitive integrin signals and ion channels in cells are necessary components in the mechanical communications[15; 16]. Another mechanistic advancement is the demonstration that cells are capable of rapidly sensing traction force in the matrix or substrate deformation, which results in directional migration and tissue-level assembly[24; 25]. It has also been proved that cells can directly sense the mechanical force from the substrate besides conventional stiffness[26]. From computational simulation along with experimental verification, COL fiber bundles re-organized by cell contraction bear the major tensile force in the ECM, which can be transmitted to distant cells[27]. During migration of epithelial cell sheet, there is also long-range force transmission through cell-cell junctions, and ERK signal wave is activated in cells to promote the migration direction[13; 28]. Cell mechanical communications also play a role in tissue pattern formation[8]. Our work has shown that cells can form stable connections through mechanical interactions on the elastic ECM, but not happening on coated glass, and the assembled connections rely on cell contraction force through the ECM substrate[24]. The cellular mechanism for distant mechanical communications and resulted cell connections still remains largely to be elucidated.

In this work, we applied the experimental model of mechanical communication from our recent study[24], and investigated the mechanosensitive pathways in regulating the mechanical attractions and cell connection stability. Our work identified the critical functions of calcium channels on the ER membranes and integrin signal along with actomyosin contraction in cell distant mechanical attractions, and N-cadherin in stabilizing cell-cell connections.

## Materials and Methods

### Cell culture and reagents

Primary airway smooth muscle (ASM) cells were originated from 6-8-week-old Sprague–Dawley rats, as described previously (approved by the Ethics Committee of Changzhou University on Studies Ethics, Grant No. NSFC 11532003)[29]. ASM cells were cultured in low-glucose DMEM (Invitrogen) supplemented with 10% FBS and penicillin/streptomycin antibiotics. Cells were maintained in the humidified culture incubator containing 5% CO2 at 37°C. The cells applied in the experiments were generally within 10 passages during regular culture.

Matrigel was purchased from BD Biotechnology, and Type I Collagen was from Advanced Biomatrix. The chemical reagents 2-Amino-ethoxydiphenyl borate (2-APB, #D9754-10G), Nifedipine (#N7634-1G), Cytochalasin D (1 μM), Blebbistatin (20 μM), and ML-7 (20 μM) were purchased from Sigma-Aldrich, and Thapsigargin (TG, #ab 120286) from Abcam. ON-TARGETplus SMARTpool N-cadherin siRNA (N-cadherin siRNA, #M-091851-01-0005) was purchased from Horizon Discovery. Integrin β1 (ITGB1) siRNA (#AM16708) were purchased from Thermo Fisher Scientific.

### Preparation of hydrogel in the Polydimethylsiloxane (PDMS) mold

The preparation processes of PDMS mold and cell network culture have been introduced in our recent work[24]. Briefly, a thin layer of PDMS (~600 μm in thickness) was generated by mixture of the two liquid components from the Sylgard 184 kit (Dow Corning). The PDMS sheet was cut into circular pieces on which one or more holes with 0.6 cm in diameter were created by a mechanical puncher. The PDMS mold was sterilized and attached onto the glass-bottom dish (NEST). Matrigel containing 0.5 mg/mL COL was added into the PDMS mold on ice, and then placed into the incubator to solidify at 37 °C for 30 min. To seed cells, about 30 μL of cell suspension was added on top of the hydrogel and stayed for 30 min in the incubator before addition of more culture medium.

### siRNA transfection and inhibitor applications

ASM cells were cultured to 40% of confluency in 6-well plates, and then transfected with 25 nM Integrin β1 or N-Cadherin siRNA, or control siRNA (final siRNA concentration in medium) by using Lipofectamine 3000 transfection kit (#L3000-008, Thermo). After 12 h, the medium was replaced with 10% FBS DMEM, and 72 h after transfection, cells were ready for experiments.

After the cells were planted on the hydrogel in the glass-bottom dish placed into 6-well plate container (Zeiss), 2 mL regular culture medium was added containing appropriate concentration of inhibitor. The experimental concentrations of 2-APB, Nifedipine, and Thapsigargin were 100 μM, 10 μM, and 10 μM, respectively.

### Time-lapse microscopy imaging

As described recently[24], the epi-microscopy system (Zeiss) was equipped with the X-Y-Z control stage for multi-position function, fine auto-focusing for time-lapse imaging, and temperature (37°C)-CO_2_ (5%) chamber to maintain cell culture conditions. Most of the imaging experiments on hydrogel were visualized with 20× objective, and the interval time was generally set as 30 min. The imaging durations were generally within 18-24 h. During the fluorescence imaging of cells stained with WGA, the excitation light from the lamp was reduced to 1/8 of the full power, and interval time was set as 1 h to minimize the photobleaching.

### Trajectory analysis of cell movements

To track cell movements, the stacked images were input into ImageJ, and the distance in pixels was transduced to length in μm on the images by the “Set Scale” function. Then by using the “Manual tracking” function from the Plugin list (Plugins ->Tracking ->Manual tracking ->Add track), the time-sequence positions (x, y) and moving distances were generated automatically by continuously clicking the target cells through the first to last frames. The acquired digital file was further input into MatLab software to calibrate the move distances and generate the map of trajectories. The movement quantifications were generally from imaging experiments within the time ranges of 18-24 h. The migration velocities and speeds of the cells were calibrated manually based on the move distances, displacements and time durations.

Graphpad and Origin2020 were applied for statistical analysis and generation of data graphs. The values on the graphs represent the mean ± S.D. (standard derivation) from their groups (in scattering dots). *, **, *** and **** indicate P < 0.05, 0.01, 0.001 and 0.0001 for significant difference between each two groups from Student’s t-test.

## Results

### Characterization of cell directional migrations during mechanical communications

Recent studies including our work have further established cell-cell distant communications through traction force sensation. To investigate the responsible cellular mechanosensitive pathways, we utilized the experimental model of cell directional migration induced by the mechanical interactions. Basically, cells were seeded on Matrigel hydrogel containing 0.5 mg/ml type I collagen (COL), and ASM cells formed branching networks in one day (Fig. 1A).

**Figure 1.**
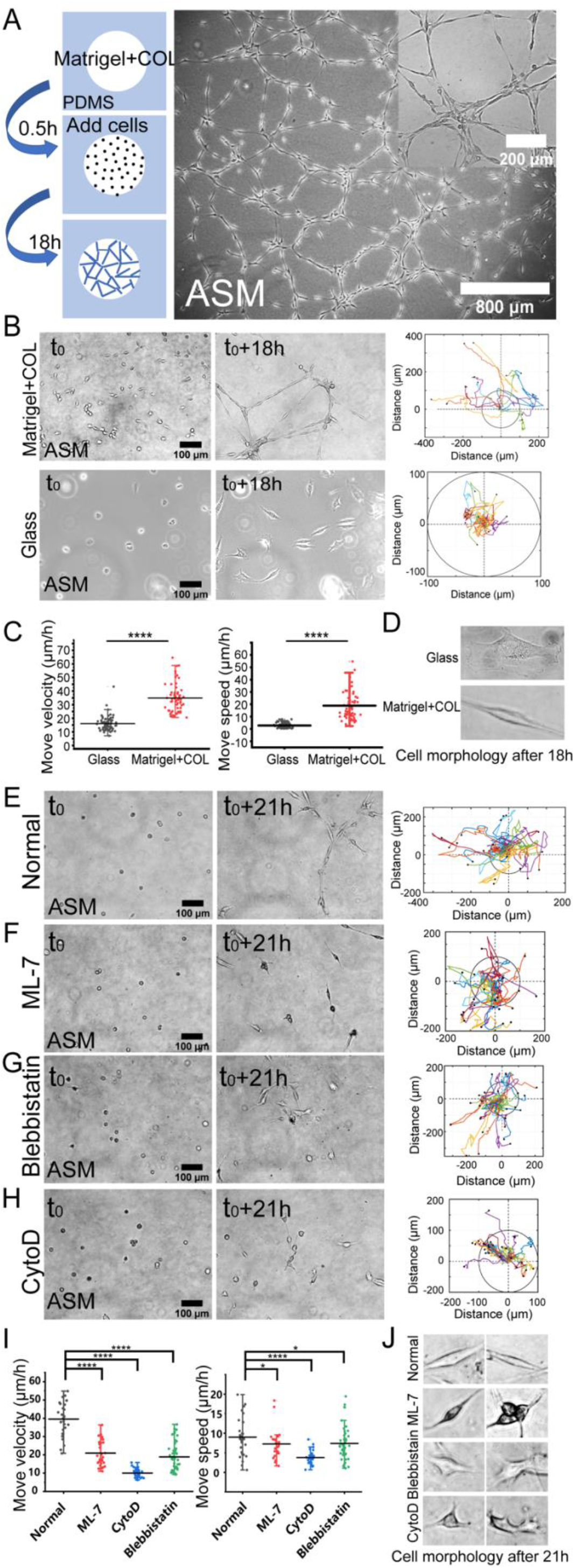
Actomyosin contraction force-mediated cell directional migrations during network assembly on the hydrogel. The hydrogel refers to 3D Matrigel containing 0.5 mg/mL COL in this work. The trajectory analysis of cell movements was done by ImageJ as described in the Methods. **(A)** Experimental setup, and the assembled network structures of ASM cells after 20-h culture. The images were taken with 5x and 20x objectives under the microscope. **(B)** Time-lapse imaging of ASM cells on the hydrogel and glass, and the trajectory analysis of cell movements. The images were taken with 20x objective for 18 h. The indicated circles present the size of 200 μm in diameter. **(C)** Statistical quantifications of cell move velocity and speed (mean ± S.D.) on the hydrogel and glass in 18 h. Move velocity (μm/h): 16.0 ±5.2, n= 72 on glass; 34.9 ± 10.4, n = 60 on hydrogel, and move speed (μm/h): 2.6 ± 2.0, n = 45 on glass; 14.5 ± 13.5, n = 32 on hydrogel. **(D)** Representative cell morphologies on the glass and hydrogel. **(E-H)** Representative cell images at 0 h and 21 h, and trajectory analysis under control condition (DMSO) (**E**), or incubation with ML-7 (20 μM) (**F**), Blebbistanin (25 μM) (**G**), and CytoD (2 μM) (**H**). **(I, J)** Statistical quantifications of cell move velocity and speed within 21 h (**I**) and representative cell morphologies (**J**) under the indicated conditions of (**E-H**). The cell sample size n = 29 (DMSO), 28 (ML-7), 31 (CytoD), 47 (Blebbistanin), respectively. Statistical comparisons were done by Student’s t-test analysis between each two groups, and *, **, ***, **** indicate P < 0.05, 0.01, 0.001, 0.0001 for significant difference, respectively, and so on through the paper.

To characterize the directional migrations, time-lapse imaging was taken every 0.5 h for 18 h, and cell positions were tracked on the hydrogel or glass for comparison. ASM cells were assembled into network structure on the hydrogel in 18 h whereas stayed individually on the glass, and cell trajectory analysis showed efficient directional movements on the hydrogel, but in random movements with far less reached spatial ranges on the glass (Fig. 1B, Movie S1). Statistical quantifications of move velocity and speed confirmed quicker migrations of ASM cells on the hydrogel than on the glass (Fig. 1C). Cells displayed elongated morphologies from traction force sensation on the hydrogel while non-directionally spreading shapes on the glass (Fig. 1D). The move speed, which reflects the direct linear distance between the initial and final positions in 18 h, shows multiple times of difference between the two conditions. The quicker movement on the hydrogel may reflect a promoting role by the cell-cell mechanical interactions.

Our previous work demonstrated that cell mutual attraction and migration rely on sensation of traction force on the hydrogel which is derived from cell contraction[24]. Here, we tried to verify the directionally migrating model in cell contraction force-dependent manner. By inhibition of actomyosin contraction with myosin II light chain kinase (MLCK) inhibitor ML-7 or myosin II ATPase activity inhibitor Blebbistanin, or disruption of actin cytoskeleton with Cytochalasin D (CytoD), cell migrations were significantly attenuated (Fig. 2E-H, Movie S2), which was further verified by statistical quantifications (Fig. 2I). Particularly without integrity of actin cytoskeleton, cell directional migration was inhibited the most (Fig. 2E&F). Due to cell migrating not always straight-forwards within 10-20 h period, the quantified speed of cell movements was generally smaller than the velocity. Hence, we applied the migration model to investigate the mechanosensitive pathways for cell-cell mechanical communication and branching assembly.

**Figure 2.**
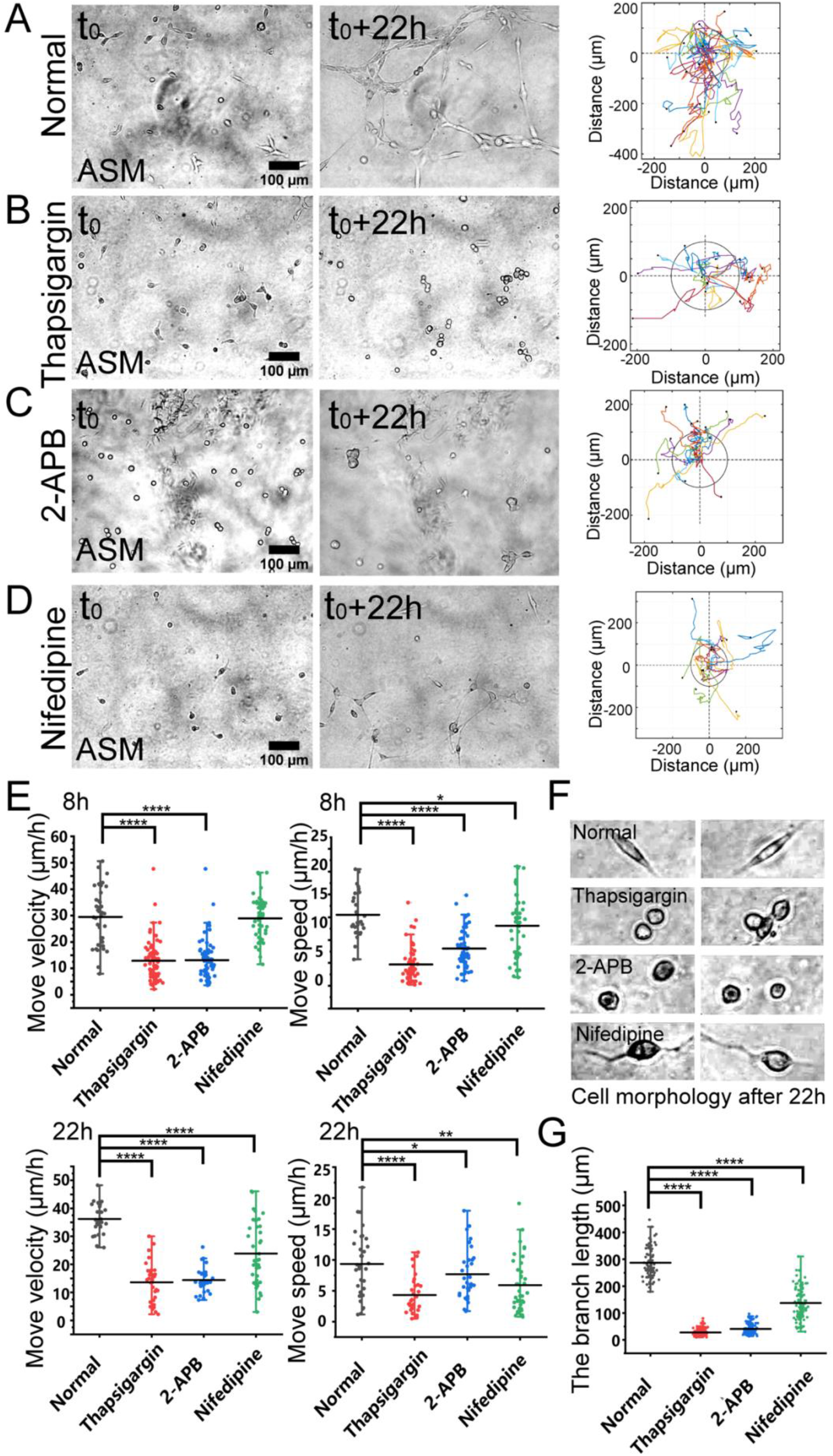
The regulations by membrane calcium channels in cell migration and branching assembly. ASM cells were seeded on the hydrogel with or without calcium channel inhibitor in the culture medium. Time-lapse imaging was taken every 0.5 h for 22 h, followed with trajectory analysis and branching quantification. **(A-D)** The cell images at 0 h and 22 h, and trajectory analysis of cell movements under the control condition (DMSO) (**A**), or with Thapsigargin (10 μM) (**B**), 2-APB (100 μM) (**C**), or Nifedipine (10 μM) (**D**) in the medium. **(E)** Statistical quantifications of cell move velocity and speed with or without the calcium channel inhibitors when culturing on the hydrogel. The sample size n = 46, 55, 34, 36 (velocity) and 56, 55, 34, 35 (speed) for 8 h (top panel), and n = 29, 28, 31, 37 (velocity) and 32, 31, 34, 40 (speed) for 22 h (bottom panel), respectively. In considering that at later time, cells might have formed branching or clusters and stopped directional migration, so quantifications were performed at 8 h and 22 h, respectively. **(F)** Representative cell morphologies with or without the calcium inhibitors. **(G)** Statistical quantifications of the branching length with or without the calcium channel inhibitors when culturing on the hydrogel for 22 h (n = 81, 112, 112, 102, respectively).

### Calcium channels regulate mechanosensation-induced cell directional migration

Based on the migration model (Fig. 1), we further investigated the cellular mechanosensing pathways. Calcium channels on the cellular membranes show mechanical sensitivity[30]. By selective inhibition of inositol 1,4,5-trisphosphate receptor (IP_3_R) calcium channel with 2-APB or the SERCA (SarcoEndoplasmic reticulum calcium ATPase) pump with Thapsigargin on endoplasmic reticulum (ER) membrane, cells showed attenuated directional migrations, and significantly inhibited branching formations (Fig. 2A-C, Movie S3). Inhibition of L-type calcium channel with Nifedipine on the plasma membrane also resulted in down-regulated cell migration and branching assembly (Fig. 2D, Movie S3). Statistical quantifications confirmed reduced migration after inhibition of these calcium channels, but less reduced migration in the early 8 h with Nifedipine treatment in comparison to the control group (Fig. 2E), suggesting early-time mechanosensation less dependent on these calcium channels. Morphologically, cells displayed round shapes with IP_3_R or SERCA inhibition, and long membrane protrusions with L-type calcium channel inhibition, in comparison to the normal elongated cell body culturing on the hydrogel (Fig. 3F). Particularly, inhibitions of the calcium channels and pump resulted in losing branching assembly (Fig. 3G). These data indicate that membrane calcium channels were mechanosensitive components during the cell-cell mechanical interactions.

**Figure 3.**
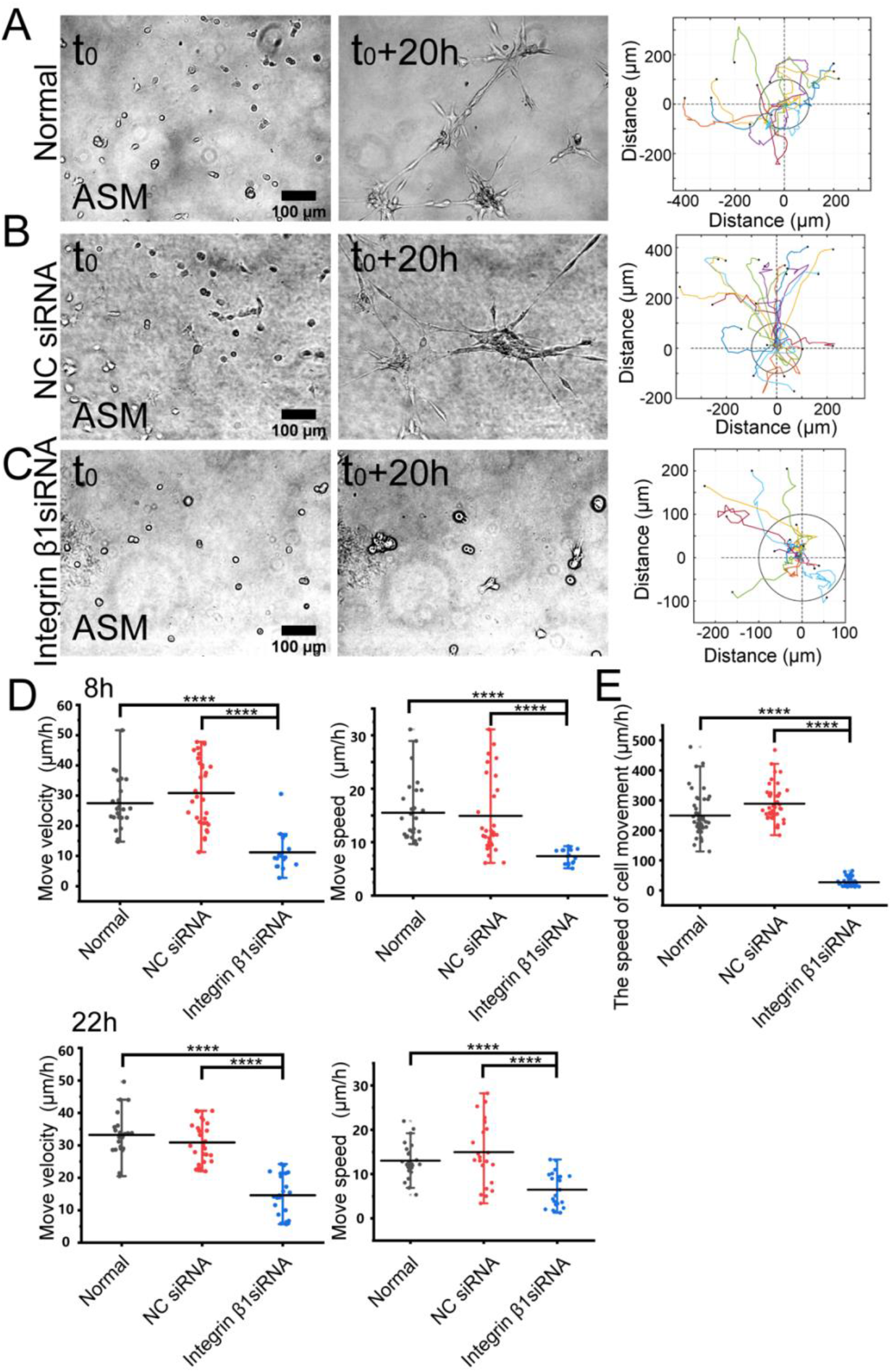
The role of integrin β1 in cell directional migration. ASM cells were transfected with control or ITGB1 siRNA, and after 72 h, cells were seeded on the hydrogel for time-lapse imaging. **(A-C)** Cell branching assembly and trajectory analysis in ASM cells at normal condition (**A**), or transfected with control siRNA (**B**) or ITGB1 siRNA (**C**). **(D, E)** Statistical quantifications of cell move velocity and speed in 8 h (top panel, n = 23, 34, 16, respectively) and 22 h (bottom panel, n = ~23, ~20, 22, respectively) (**D**), and assembled branching length (n = 44, 44, 48, respectively) (**E**) under the indicated conditions.

### Integrin β1 mediates cell directional migration and branching assembly

Integrin β1 forms heterodimers with variable integrin α subunits (α_1-11_, or α_v_)[31], so we tried to inhibit β1 subunit and checked the role in mechanosensation-induced cell directional migration. After transfected with Integrin β1 (ITGB1) siRNA, cell directional migration was significantly reduced in comparison with normal condition or control siRNA (Fig. 3A-C, Movie S4). Especially, the branching structure couldn’t be assembled in ASM cells transfected with ITGB1 siRNA (Fig. 3C). As shown by further quantifications, ITGB1 siRNA transfection resulted in inhibited cell directional migration (Fig. 3D), and nearly blocked branching assembly (Fig. 3E). These results indicate that integrin α_(x)_ β1 is the mechanosensitive component for the cell-cell mutual distant interactions.

### N-cadherin regulates cell branching formation

N-cadherin is one mechanosensitive molecule on the plasma membrane and regulates cell-cell junctional connections[32]. We further investigated the role of N-cadherin in cell migration and branching assembly during the cell-cell mechanical interactions. ASM cells displayed directional migration and branching assembly under normal condition or with control siRNA, and showed regular migration and formed cell clusters instead of branching structures with N-cadherin siRNA (Fig. 4A-C, Movie S5). Statistical quantifications demonstrated that ASM cells maintained regular directional migrations after N-cadherin siRNA transfection, but failed in assembly of branching structures (Fig. 4D). From the quantifications (Fig. 4D), the cells seemed moving more with N-cadherin siRNA, which might be due to contraction of the cell groups, whereas cells were more stabilized in the branches under the control conditions. These results indicate that N-cadherin wasn’t necessary for cell distant mechanosensation, but essential for cell-cell junctional connections to form stable branching structures.

**Figure 4.**
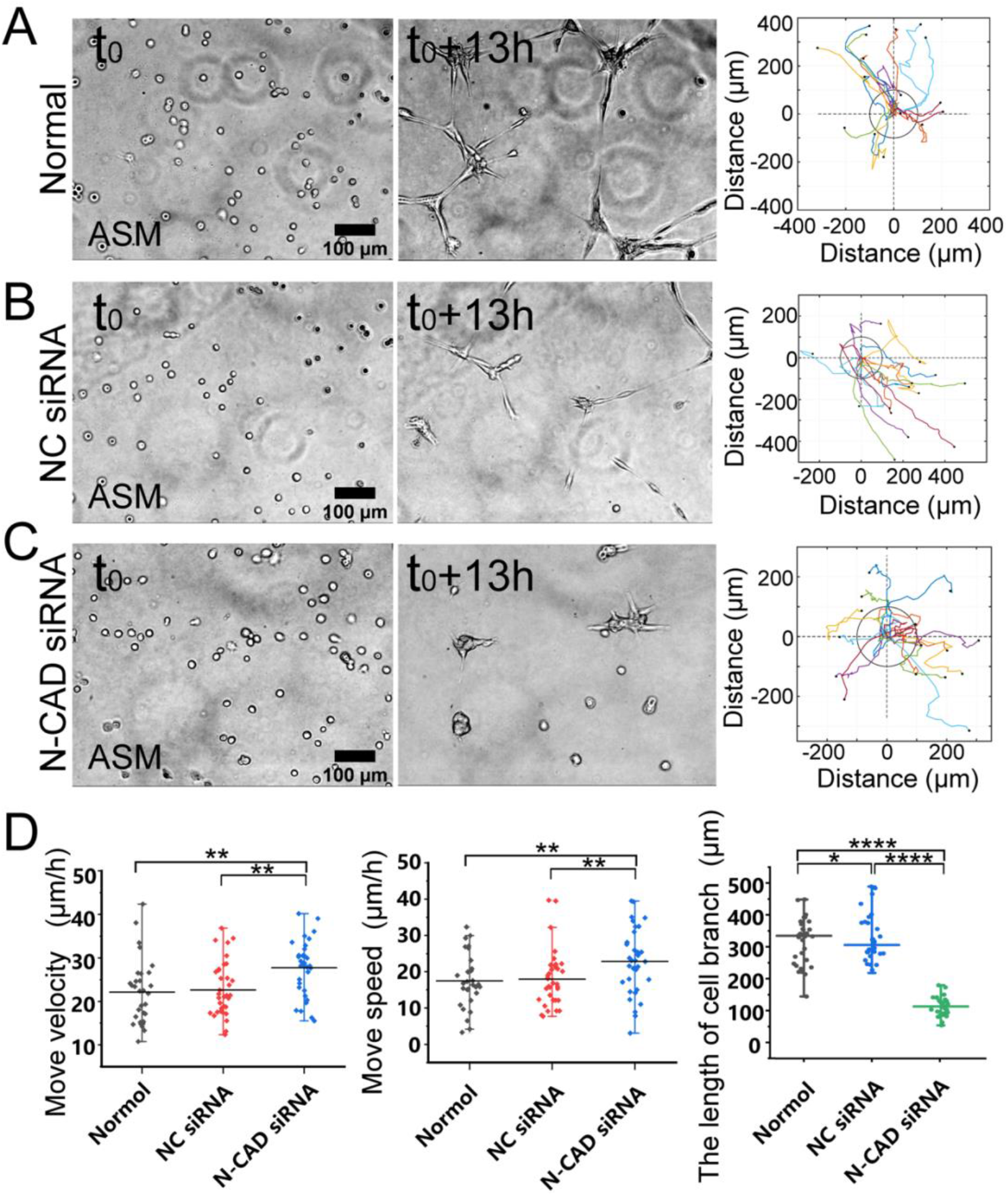
N-cadherin in regulating cell branching formation. ASM cells were transfected with N-cadherin (N-CAD) or control siRNA, and after 72 h, cells were seeded on the hydrogel for time-lapse imaging. **(A-C)** ASM cell branch assembly and trajectory analysis at normal condition (**A**), or transfected with control siRNA (**B**) or N-CAD siRNA (**C**). Cells transfected with N-CAD siRNA formed clusters without further migration later, and the first 13-hour images were analyzed. **(D)** Statistical quantifications of ASM cell move velocity and speed (n = 33, 34, 37, respectively) as well as length of assembled branches (n = 29, 29, 38, respectively) at the indicated conditions of (**A-C**).

## Discussion

Cell-cell mechanical communications in long or large spatial scale have been increasingly recognized in the past decade. It has been further revealed recently that cells are able to communicate via traction force mechanosensation through the matrix substrates. The mechano-responsive mechanism is still to be investigated. In this work, we tried to study the primary mechanosensing components for cell-cell distant mechanical interactions. Based on the experimental model from our recent work[24], we characterized cell directional migration and network assembly to measure cell-cell mechanical attractions, which was applied to identify cellular mechanosensitive components.

ASM cells assembled into network structures within one day on the hydrogel, while stayed isolated individually on glass surface without force transmission (Fig. 1A-D, Movie S1). Migration quantifications showed much quicker movements on the hydrogel than on glass, indicating a significant promoting role in driving cell directional migration by cell-cell mechanical interactions (Fig. 1C). The integrity of actin cytoskeleton and actomyosin activity are essential for the mechanical interactions (Fig. 1E-I, Movie S2), verifying the interaction force originated from cellular contraction. Hence, we have validated the experimental model based on migration and branching assembly for cell-cell mechanical communications.

Recent studies reported that Piezo ion channel (one type of mechanosensitive calcium channel) on the plasma membrane plays an important role in cell mechanosensation of force[15; 16]. Here, we further explored the relevance of calcium channels on plasma and ER membranes during the cell-cell distant mechanical interactions. The experimental data demonstrated that the IP_3_R calcium channel and SERCA pump on ER membrane are important for cell directional migration and essential for branching assembly (Fig. 2A-E, Movie S3). This observation is consistent with the previous reports that calcium signal from ER storage is mechanosensitive to mechanical stimulation[30; 33], which also supports the cell-cell distant attractions due to mechanical communications.

Integrins are mechanosensitive receptors on the plasma membrane in response to ECM mechanics[34; 35]. Two recent studies have shown that integrin signaling mediates intracellular mechanical interactions[15; 16]. Here, we tried to validate the role of integrin signaling in this experimental model. Inhibition of integrin β1 with siRNA in ASM cells, which is a representative subunit forming dimer with variable α subunits, led to significantly reduced cell-cell remote attractions as shown by the inhibited directional migrations and branching formations (Fig. 3A-E, Movie S4). In considering that integrins are basic components in assembling focal adhesions[36], deficiency in integrin may hinder the mechano-response for directional migrations towards the neighboring cells. From these few studies, it is conclusive that integrins are mechanosensitive signals in the cell-cell distant mechanical communications.

There is an interesting observation in our work that ASM cells formed one-to-one connections to assemble continuous network structure, but no stable connections occurred on glass without traction force transmission (Fig. 1A)[24]. N-cadherin is one mechanosensitive molecule at cell-cell junctions[32], and also shows the role in promoting cell directional movements[37]. In this work, selective knockdown of N-cadherin with siRNA didn’t have apparent impact on cell-cell remote attraction as measured by the move speeds, but inhibited one-to-one stable cell connections for branching assembly (Fig. 4A-D, Movie S5). This result implies that N-cadherin is critical for traction force-induced stable cell-cell connections.

In summary, our work identified the importance of IP_3_R calcium channel and SERCA pump on the ER membrane and also verified integrin signal in the cell-cell distant mechano-attractions, and further demonstrated junctional N-cadherin in stabilizing traction force-induced cell-cell connections. The molecular mechanism or subsequential relations of these components during the mechanosenation need more studies.

## Acknowledgements

This work was supported financially by National Natural Science Foundation of China (NSFC 11872129), Natural Science Foundation of Jiangsu Province (BK20181416), and Jiangsu Provincial Department of Education (M.O.); National Natural Science Foundation of China (11532003) (L.D.); National Natural Science Foundation of China (11902051) (B.B.).

## Author contributions

M. Ouyang and L. Deng conceived the project and designed the research; Y. Zhu performed the majority of experiments; Y. Zhu and M. Ouyang did the major data analysis & organization; J. Wang and Q. Zhang helped with experiments and data analysis; J. Guo, and B. Bu joined in project discussion and provided technique supports; L. Deng provided the setups of equipment; M. Ouyang, L. Deng, and Y. Zhu prepared the paper.

